# Combining optical imaging of cleared tissue with mathematical modelling to predict drug delivery and therapeutic response

**DOI:** 10.1101/219865

**Authors:** Angela d’Esposito, Paul Sweeney, Rebecca Shipley, Simon Walker-Samuel

## Abstract

Understanding how drugs are delivered to diseased tissue, and their subsequent spatial and temporal distribution, is a key factor in the development of effective, targeted therapies. However, the interaction between the pathophysiology of diseased tissue and individual therapeutic agents can be complex, and can vary significantly between individuals. In cancer, suboptimal dosing resulting from poor delivery can cause reduced treatment efficacy, upregulation of resistance mechanisms and can even stimulate growth. Preclinical tools to better understand drug delivery are therefore urgently required, which incorporate the inherent variability and heterogeneity of human disease. To meet this need, we have combined multiscale mathematical modelling, high-resolution optical imaging of intact, optically-cleared tumour tissue from animal models, and in vivo magnetic resonance imaging (MRI). Our framework, named REANIMATE (REAlistic Numerical Image-based Modelling of biologicAl Tissue substratEs) allows large tissue samples to be investigated as if it were a living sample, in detailed, highly controlled, computational experiments. Specifically, we show that REANIMATE can be used to predict the heterogeneous delivery of specific therapeutic agents, in disparate two murine xenograft models of human colorectal carcinoma. Given the wide adoption of optical clearing equipment in biomedical research laboratories, REANIMATE enables a new paradigm in cancer drug development, which could also be applied to other disease areas.

## Introduction

Mathematical modelling of biological tissue is increasingly used to better understand complex biological phenomena, and to better understand the development of disease.^1^ This developing paradigm of computational experiments enables complex and subtle interventions can be performed in a manner that would be challenging or imposible in a conventional experimental setting. Multiple hypotheses can be iteratively generated and tested, allowing new research avenues to be discovered, and for them to be rapidly and cheaply evaluated, at scale.

In this study we present a novel framework for performing realistic computational experiments that naturally incorporates the variability and heterogeneity found between biological samples (even those derived from the same disease process). It allows large tissue samples to be imaged and treated as living specimens, by combining cutting-edge optical imaging techniques with mathematical modelling. We have named our framework REANIMATE (REAlistic Numerical Image-based Modelling of biologicAl Tissue substratEs) (see **Figure 1**).

**Figure 1.**
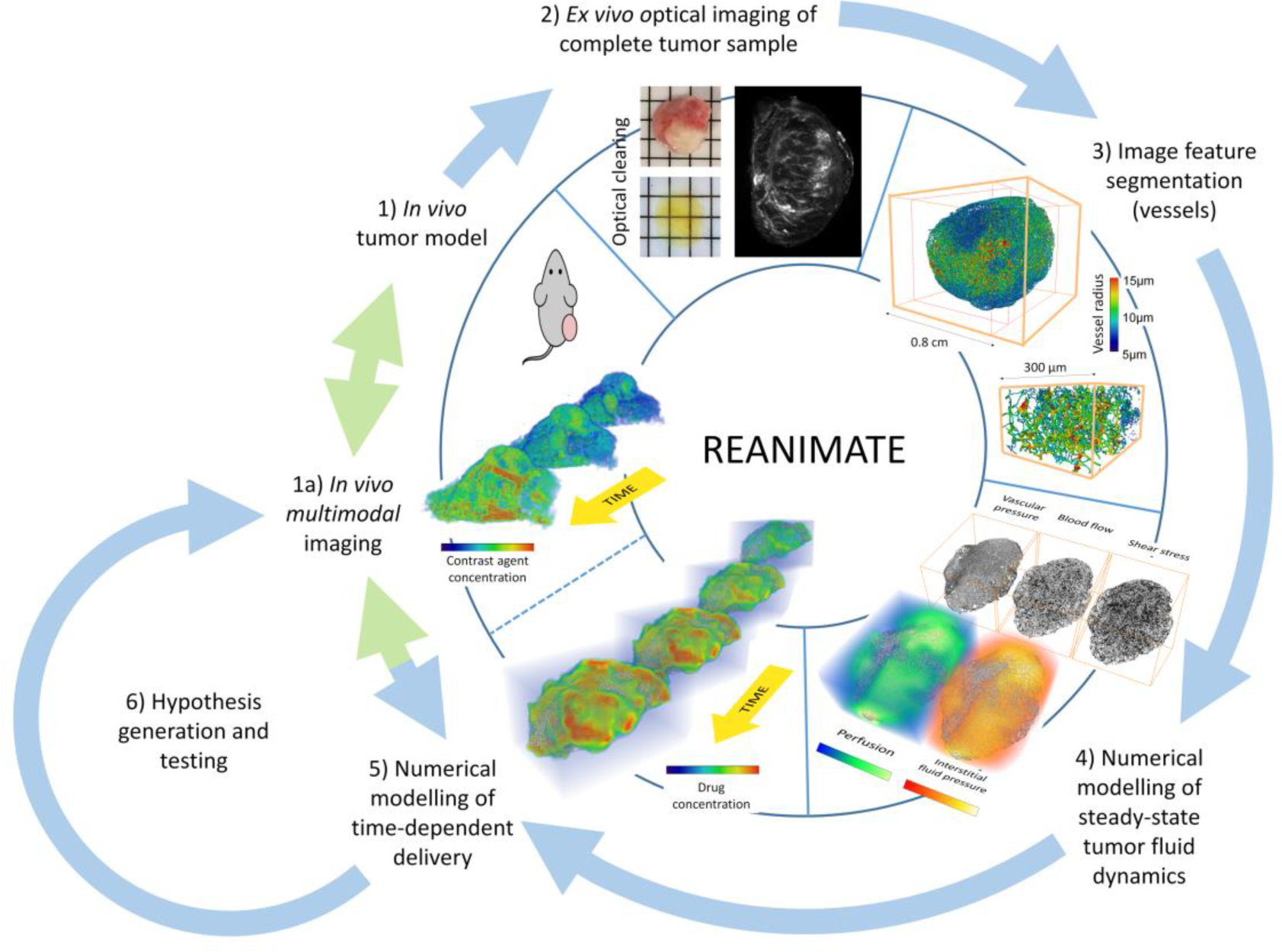
The REANIMATE pipeline for *in vivo* and *ex vivo* imaging of intact tumours and performing three-dimensional computational fluid mechanics simulations. After *in* vivo imaging (1), which can be performed longitudinally during tumor growth, tumours are resected and optically cleared (2), to render tumours transparent for three-dimensional fluorescence imaging. Optical images are processed to segment fluorescently-labelled structures within the tumor microenvironment (3) (in this case, blood vessel networks), which are reconstructed in 3D graph format (nodes and connecting segments, each with a radius corresponding to the size of the blood vessel). These geometrical data become the substrate for computational fluid dynamic models to estimate steady-state blood flow and interstitial transport (4) and time-dependent numerical modelling of drug delivery (5). All of these data can then be used to perform *in silico* experiments (e.g. assessing the heterogeneous delivery of drugs or contrast agents), which can be compared with *in vivo* experiments in the same tumor models, or even the same mice. In this study, REANIMATE is used to study the action of a vascular disrupting agent (Oxi4503) in two models of human colorectal carcinoma.

For the results generated by computational experiments to be confidently accepted, careful experimental validation must be performed. Through its use of optical image data from cleared (transparent) tissue, REANIMATE can be used to perform computational experiments on realistic, large,^2^ high resolution structural data, rather than relying on small, isolated samples or synthetically-generated substrates. Optical imaging of cleared tissue can provide spatial data on complex, interacting structures (such as blood vessel networks, cell nuclei, etc.), which can be explored, across entire organs,^3,4^ and at resolutions of a few microns,^2^ by administering fluorescently-labelled probes that bind to specific strutures.

Capturing the physiological variation in complete, intact tissue specimens is particularly useful in tumours, which can be highly heterogeneous, both between tumour types, tumour deposits and even within individual tumors.^5^ This results in substantial differences in, for example, drug delivery, oxygenation and gene expression,^6^ with associated differences in therapeutic response and resistance. Effective therapy normally requires drugs to be delivered to the site of disease, at as high a concentration as possible, but avoiding signficant toxicity effects in healthy tissues, whilst sub-optimal exposure can limit treatment efficacy, induce exposure-mediated resistance mechanisms,^7^ or even stimulate tumor growth.^8^

This complex physiological-pharmacological landscape requires careful analysis in order to be fully understood, and is most appropriately and naturally approached within a mathematical modelling framework. The numerical modelling component of REANIMATE consists of two steps: first, a solution is sought from a set of coupled fluid dynamics models that describe steady-state vascular and interstitial fluid transport; second, the steady-state solution (or set of solutions) is used to parameterise a time-dependent model that describes the vascular and interstitial uptake of exogenousy administered material. This can be used, for example, to model the heterogenous pharmacokinetics of drug or imaging contrast agents, or delivery of individual particles (e.g. T-cells, antibodies). Terms can also be introduced to describe drug targetting and metabolism.

As a first evaluation, we have used REANIMATE to: 1) study the spatially heterogeneous uptake of a gadolinium-based MRI compound (which allowed us to compare nuerical modelling solutions with ground-truth *in vivo* data); and 2) investigate the effect of the vascular disrupting agent (VDA) Oxi4503 on tumour vasculature. These results provided a rich, three-dimensional framework for probing spatially heterogeneous tumour drug delivery and treatment response.

## Results

### Preparation of tissue substrates for mathematical modelling from optically-cleared tissue samples

SW1222 and LS174T human colorectal carcinoma tumours have been extensively studied, by our group and others, with SW1222 tumors displaying greater cell differentiation, more uniform vasculature and greater perfusion than LS174T tumors.^9-15^ These tumours formed the basis of our initial evaluation of the REANIMATE framework. Mice bearing each tumour type (n=5 of each) were grown subcutaneously, for 10 to 14 days, to a mean volume of 83 ± 12 mm^3^. These mice were injected intravenously with fluorescently-labelled lectin (AlexaFluor-647) and, following a circulation time of 5 minutes, tumors were resected, optically cleared with BABB, and imaged with optical projection tomography (OPT) ^16^.

Blood vessels were segmented from OPT images using Frangi filtering ^17^ and thresholding, then converted into graph format, consisting of nodes (branch points) and vessel segments. These networks were typically composed of 30,000 to 200,000 nodes (**Figure 2a**). We performed a comparison of the vessel architecture from each tumor type with previously published data, which showed that vessel architecture was preserved during tissue clearing (see **Supplemental Materials**).

**Figure 2.**
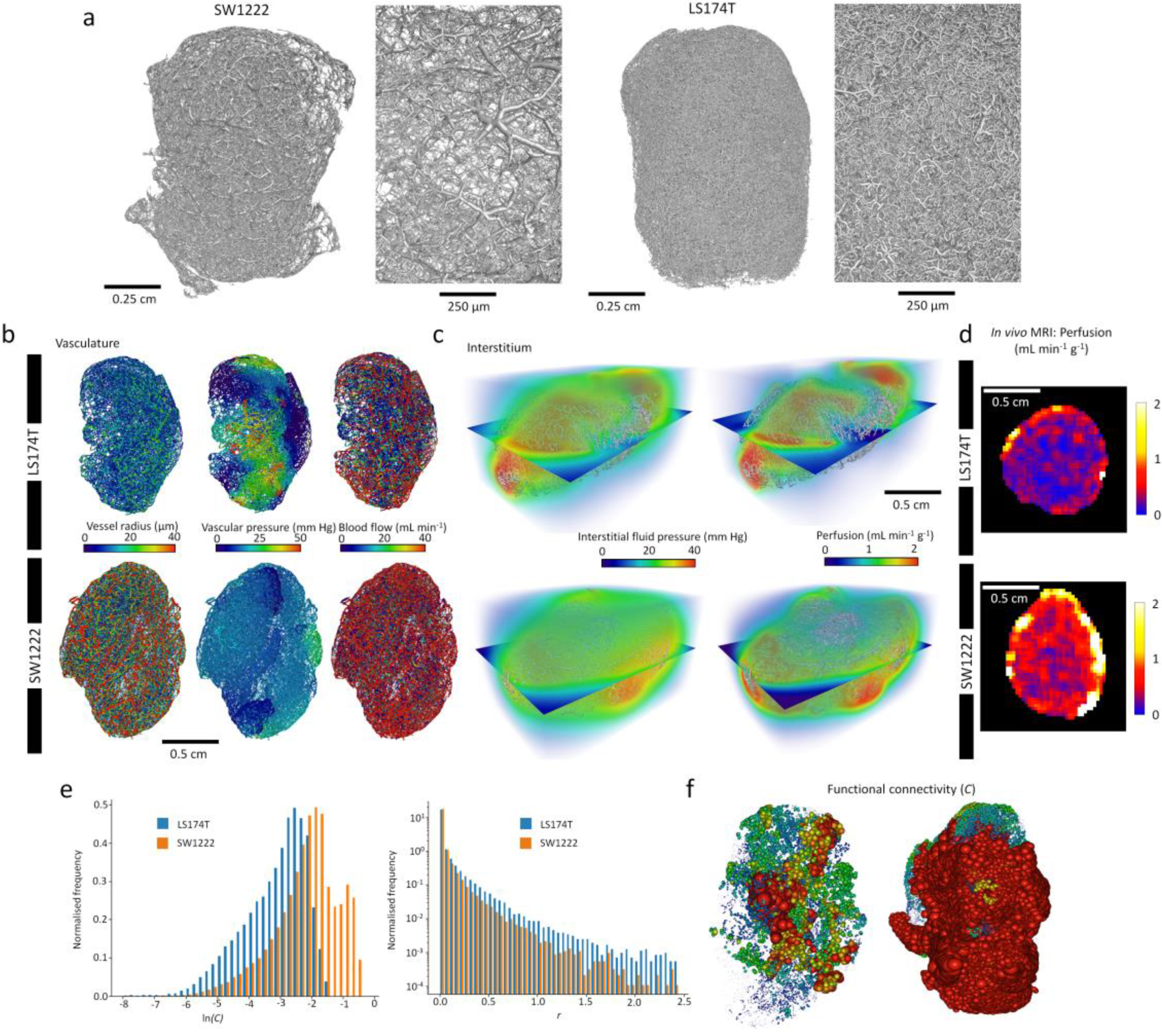
Summary of blood vessel geometry measures (vessel diameter, inter-branch distance, branching angle and inter-vessel distance), steady-state fluid dynamics simulation parameters (vascular pressure, blood flow, interstitial fluid pressure, perfusion), and whole-vascular network functional connectivity and redundancy measures (node connectivity and redundancy distance). These were taken from blood vessel networks reconstructed from LS174T and SW1222 xenografts. (a) Reconstructions of the segmented networks (diameters scaled according to their measured values), demonstrating clear differences in vascular architecture between the tumour types, as well as intra-tumour spatial heterogeneity. (b) Vascular networks colored according to vessel radius, simulated blood flow and intravascular pressure in two example colorectal carcinoma xenografts (LS174T and SW1222), derived from whole-tumour optical image data. Also shown are the same vascular networks overlaid with simulated interstitial fluid pressure and perfusion values. (c) Example *in vivo* measurements of tumor perfusion, acquired using arterial spin labelling MRI, in LS174T and SW1222 tumors. (d) Frequency distributions of ln(node connectivity) and redundancy distance ratio, demonstrating clear distinctions in the distributions for the two tumour types. (e) Tumor vessel networks, with nodes scaled according to vessel connectivity; the larger the node, the greater the connectivity.

### REANIMATE simulation of steady-state tumor blood flow and tissue perfusion

Our first aim was to use these whole-tumor blood vessel networks as the substrate for novel simulations of steady-state, whole-tumor fluid dynamics. Our steady-state mathematical model of tumour fluid dynamics was comprised of coupled intravascular and interstitial compartments, with exchange mediated by vascular permeability. Blood flow and interstitial delivery were modelled using Poiseille flow and Fick’s law, respectively, and we optimised our steady-state model over the entire tumor by assigning semi-stochastic pressure boundary conditions to inlet vessels. We performed our initial simulations on six colorectal tumor xenografts (n=3 SW1222 and n=3 LS174T).

Solutions to our mathematical model predicted significant differences in blood flow, blood velocity and vessel wall sheer stress, between the SW1222 and LS174T tumour types, which is consistent with their known characteristics (see **Figure 3**, **Table 1**). Key to the interpretation of these results was our ability to compare them directly with equivalent *in vivo* imaging data (in this case arterial spin labelling magnetic resonance imaging (ASL MRI)), which can be used to quantify perfusion, noninvasively ^9^. Perfusion is a measure of the rate of delivery of fluid to biological tissue, and is dependent on blood flow, vascular permeability and interstitial density, amongst other factors. We regridded simulated perfusion values to the resolution of *in vivo* images (150 μm) for direct comparison. Both *in vivo* measurements and simulations showed that SW1222 tumours were better perfused than LS174T tumors, which is again consistent with the results of previous studies.^18^

**Figure 3.**
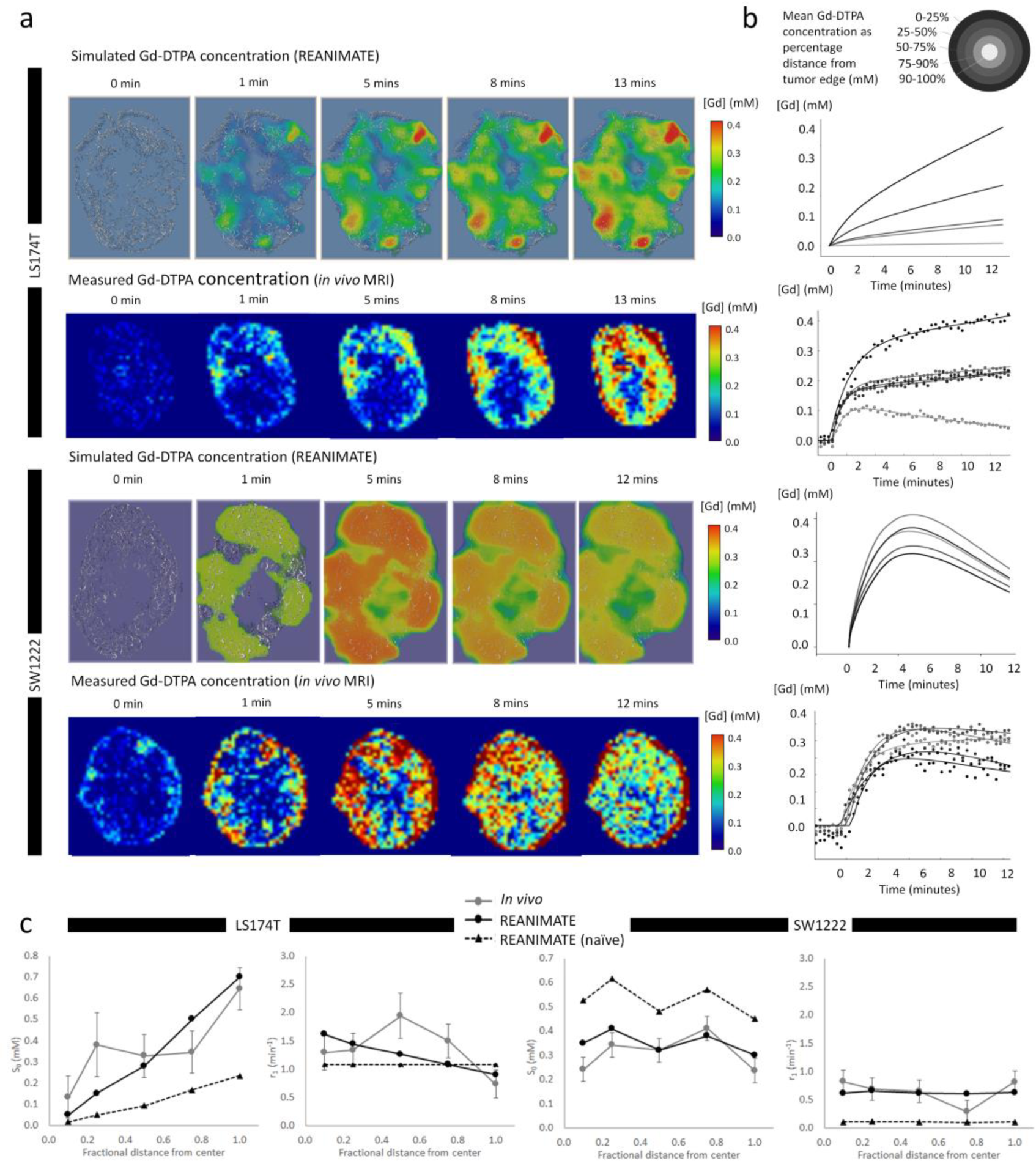
REANIMATE simulation of Gd-DTPA (an MRI contrast agent) delivery to LS174T and SW1222 tumor, guided by *in vivo* data. (a) Spatial distributions of Gd-DTPA concentration in each tumor type, with example *in vivo* data shown below. (b) Mean Gd-DTPA concentration as a function of time, for REANIMATE and *in vivo* data. Each curve shows the average uptake at a fractional distance between the perimeter and centre of mass of the tumor. (c) Pharmacokinetic analysis of Gd-DTPA uptake curves. All Gd-DTPA concentration ([Gd]) data were fitted to a phenomenological model of the form 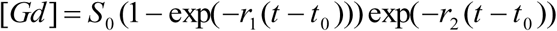 where *S*_0_ represents the peak concentration and *r*_1_ and *r*_2_ the wash-in and washout rates, respectively. Plots show the mean value (± standard error) for *S*_0_ and *r*_1_ parameters, as a function of fractional distance from the tumor periphery. REANIMATE data are shown for naïve analysis (i.e. using fixed, literature parameter values) (dashed lines) and corrected by iteratively fitting REANIMATE data to *in vivo* data.

**Table 1.** Blood vessel network geometry summary statistics from optically-cleared colorectal carcinoma xenograft models (mean ± standard error). Reference values are previously published values taken from the literature, in the same tumor models (^1^ Folarin et al, 2010 ^11^).

| Parameter | LS174T | Reference | SW1222 | Reference |
| --- | --- | --- | --- | --- |
| Vessel diameter | $20 \pm 2 \mu\text{m}$ | $19 \mu\text{m}^1$ | $34 \pm 5 \mu\text{m}$ | $36 \mu\text{m}^1$ |
| Inter-branch distance | $61 \pm 8 \mu\text{m}$ | $50 \mu\text{m}^1$ | $69 \pm 9 \mu\text{m}$ | $63 \mu\text{m}^1$ |
| Inter-vessel distance | $40 \pm 4 \mu\text{m}$ | $39 \mu\text{m}^1$ | $28 \pm 5 \mu\text{m}$ | $79 \mu\text{m}^1$ |
| Branching angle | $73.0 \pm 0.8^\circ$ | $78^\circ^1$ | $80.1 \pm 0.8^\circ$ | $82^\circ^1$ |
| Number of branches per node | $2.7 \pm 0.2$ | N/A | $3.4 \pm 0.3$ | N/A |
| Blood volume fraction | $0.13 \pm 0.01$ | $0.9^1$ | $0.24 \pm 0.04$ | $0.25^1$ |
| Tumour volume | $0.13 \pm 0.01 \text{ cm}^3$ | N/A | $0.12 \pm 0.02 \text{ cm}^3$ | N/A |

We found that, in both *in vivo* measurements and simulations, perfusion was distributed heterogeneously throughout the tumors, with markedly raised values at the periphery of both tumor types; however, perfusion at the centre of SW1222 tumors was much greater than in LS174T tumors, both in simulations and *in vivo* data (**Figure 2b**, **c** and **d**). On average, we found that simulated perfusion values in LS174T tumors matched those measured *in vivo*, whilst SW1222 were slightly larger, but of the same order. Arterial spin labelling estimated mean perfusion as 33 ± 18 mL min^-1^ 100g^-1^ for SW1222 tumors and 19 ± 8 mL min^-1^ 100g^-1^ for LS174T, compared with 73 ± 3 and 18 ± 7 mL min^-1^ 100g^-1^, respectively, in REANIMATE simulations.

These results show that mathematical modelling of fluid dynamics, using cleared tumor tissue as a substrate, is both feasible and provides quantitative predictions of vascular perfusion that are in keeping with experimental results. Our next step was to use the steady-state to parameterise a time-dependent model to simulate the delivery of exogenously-administered material.

### REANIMATE simulation of MRI contrast agent (Gd-DTPA) delivery

As a first evaluation, we chose to model the dynamics of chelated gadolinium (Gd-DTPA), a widely used MRI contrast agent with well-studied pharmacokinetics, and directly compare them with *in vivo* measurements. Gd-DTPA can be thought of as a proxy for a non-metabolised therapeutic agent. Time-dependent simulations were calculated using a ‘propagating front’ algorithm, that used steady-state solutions to mimic the physical delivery of material, both via vascular flow and by endothelial and interstitial diffusion.

To provide ground-truth data for comparison, and to generate new modelling substrates, we performed *in vivo* experiments to measure the delivery of a bolus of Gd-DTPA in a set of LS174T (n=5) and SW1222 (n=6) tumours, using a dynamic contrast-enhanced (DCE) MRI sequence (**Figure 3**). Following these measurements, mice were culled via cervical dislocation. We resected and set two tumours aside (one LS174T and one SW1222) for processing within the REANIMATE framework.

Here, steady-state simulations were performed as described above, and were used as the basis for time-dependent delivery simulations. The influence of each vessel network inlet was modelled independently, and an algorithm was developed that monitored a propagating front through the network. Exchange between the vascular and interstitium was cast in a finite element framework, with vessel permeability initially fixed at 1×10^-6^ cm s^-1^.^19,20^ The interstitium was modelled as a continuum with a constant cell volume fraction (*f*_c_ = 0.8 ^21^). Gd-DTPA does not cross the cell membrane,^22^ and so Gd-DTPA concentration ([Gd]) was scaled by the fractional volume of the extra-cellular space, and we assumed a constant rate of diffusion through the interstitium (*D* = 2.08×10^-4^ mm^2^ s^-1^ ^23^). Both *in vivo* measurements and simulations had a duration of 12 minutes, with a temporal resolution of 16 seconds. Simulations were driven by a bi-exponential vascular input function, taken from the literature.^24^ Initially, 1% of the entire dose of the input function was partitioned across all tumor inlets, weighted by the inflow rate for the individual inlet.

Intravascular and interstitial delivery were rendered as four-dimensional visualisations (see **Supplemental Movie 1** and **Supplemental Move 2**). Analysis of our experimental MRI measurements from LS174T tumors revealed a prolonged, peripheral enhancement pattern, which is typical of the tumor type ^18^ (**Figure 3a**). Plotting contrast agent uptake as a function of distance from the tumor edge revealed a highly heterogeneous enhancement pattern, with decreasing concentration for increasing proximity to the tumour centre (**Figure 3b**). Conversely, we found that SW1222 tumors enhanced with Gd-DTPA much more rapidly and homogeneously, with a peak enhancement at around 4 minutes, followed by a washout phase. We fitted our experimental Gd-DTPA concentration-time curves to a phenomenological model of contrast agent enhancement (Equ. 14), in which the parameter S0 defines the peak enhancement, and *r*_1_ and *r*_2_ are rates of concentration increase and decrease, respectively.^25^ Again, plotting these parameters as a function of distance from the edge of each tumor type (**Figure 3c**) confirmed our observations of decreasing contrast uptake with distance from the tumor edge in LS174T tumors, compared with a more homogeneous enhancement pattern in SW1222 tumors.

### Iterative optimisation of time-dependent delivery simulations guided by experimental results

REANIMATE solutions describing Gd-DTPA delivery were analysed in the same manner as experimental data, i.e. as a function of distance from the tumor periphery. Simulations were performed in two stages: the first estimated Gd-DTPA enhancement using the initialisation parameter values defined above (the naïve solution); the second stage used modified parameter values, based on iteratively minimising the disparity between simulated and *in vivo* data.

Our naïve analysis underestimated the magnitude of contrast enhancement in LS174T tumors, but still reflected their spatial heterogeneity, with the *S*_0_ parameter decreasing with distance from the tumor periphery. The enhancement rate parameter, *r*_1_, provided a good fit to *in vivo* data, but did not reflect its increasing value at the tumour centre. To account for this, we increased the mean vascular permeability to 0.9×10^-6^ cm s^-1^ at the periphery, with a linear increase to 1.1×10^-6^ cm s^-1^ in the centre. This provided a much better accordance with *in vivo* data (**Figure 3c**). For the SW1222 tumor naïve simulation, contrast agent uptake was overestimated, but homogeneously distributed, reflecting what was found *in vivo*. The rate of enhancement was also much greater than *in* vivo. We therefore uniformly decreased vascular permeability to 0.75×10^-7^ cm s^-1^ in the second simulation, which then provided a good accordance with *in vivo* data.

We can therefore conclude from these experiments that REANIMATE can provide good order-of-magnitude estimates of the delivery of Gd-DTPA, but which can be further finessed by *in vivo* measurements.

### Dual-fluorophore optical imaging of Oxi4503 response

Using our optimised Gd-DTPA delivery data, we then went on to investigate the ability of REANIMATE to model drug uptake and response to treatment. This required the development of a novel dual-fluorophore imaging technique that allowed measurements of tumor vascular structure at two separate time points to be encoded. We chose to model vascular targeting therapy, due to its rapid, well-characterised mechanism of action, which can be captured with *in vivo* MRI ^26^. The acute effects of VDAs have been well-documented, using histology,^27,28^ MRI ^29^ and *in vivo* confocal microscopy,^30^ which demonstrate rapid vascular shutdown and extensive vessel fragmentation within the first 60 minutes to 24 hours of administration. This causes decreased perfusion, especially in the central part of the tumour,^29,31^ and an associated increase in hypoxia and cell death.^31^ In this study, we investigated a single dose of Oxi4503, at 40 mg kg^-1^.

Our dual-fluorophore method allowed us to characterise blood vessel structure at two separate time points, by administering fluorescently-labelled lectin (AlexaFluor-568) just prior to injecting Oxi4503, and then a second lectin 90 minutes later (AlexaFluor-647). Our rationale was that vessels occluded by Oxi4503, and were no longer perfused, would be labelled by only the first fluorophore; vessels that remained perfused following therapy would be labelled with both fluorophores. As a validation of our results, *in vivo* arterial spin labelling (ASL) MRI was also performed on a subset of tumors (n=3 of each tumor type). Mice, each bearing an LS174T or SW1222 tumor, were randomly assigned to treatment (Oxi4503, 40 mg kg^-1^) or control groups (administered saline). We produced volume renderings of vessel networks in which vessels were colored blue if co-labelled with both fluorescent lectins (i.e. were perfused pre- and post-Oxi4503) at 90 minutes or green otherwise (**Figure 4a**, **Supplemental Movie 3**).

**Figure 4.**
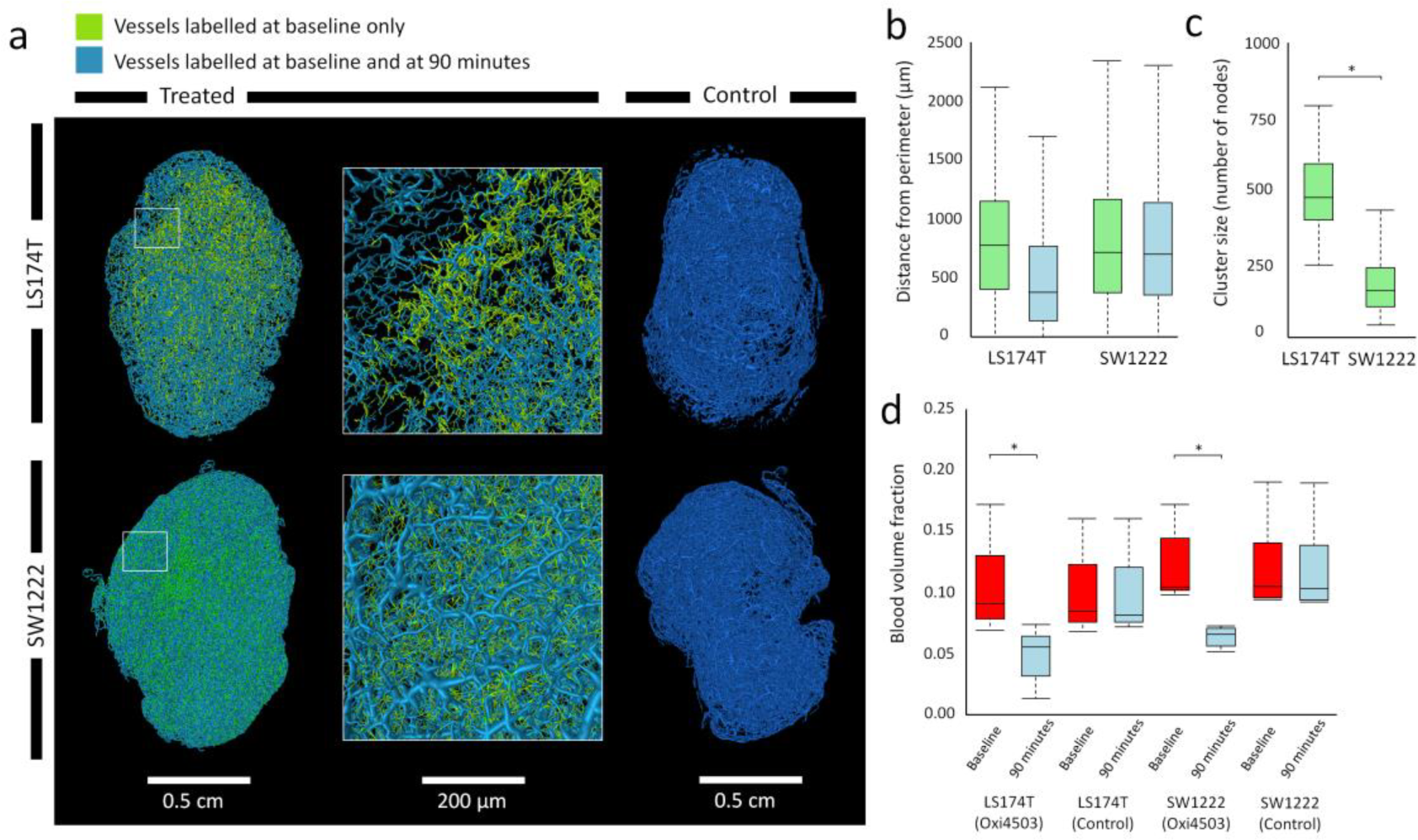
Dual-fluorophore, optical imaging of the response of colorectal carcinoma models (LS174T and SW1222) to treatment with a vascular disrupting agent (Oxi4503), at baseline and 90 minutes post-dosing. Tumors were injected with lectin labelled with the first fluorophore (AlexaFluor-568) prior to administration of 40 mg kg^-1^ of Oxi4503, to label all blood vessels in the tumors. 60 minutes later, to assess the acute and heterogeneous effects of Oxi4503, a second lectin labelled with a different fluorophore (AlexaFluor-647) was injected, to label vessels that remained perfused. (a) Whole-tumor blood vessel networks, colored according to whether they remained labelled following Oxi4503 (dual-labelled, blue) or were no longer perfused (single-labelled, green). OPT signal intensity images are also shown. (b) Box plot showing the distance of single- and dual-labelled vessels from the tumor periphery and (c) the size of single-labelled clusters in both tumor types. (a-c) show that LS174T tumors lost large vascular regions at their centre, whereas SW1222 tumors showed a more distributed pattern of perfusion loss, distributed throughout the tumor. (d) Box plot of blood volume measurements from dual-labelled Oxi4503 and control-treated tumors. Blood volume significantly reduced (p<0.001) in both SW1222 and LS174T tumors when treated with Oxi4503, whereas there was no significant difference in control tumors.

Our initial analysis of our dual-fluorophore data focussed on geometrical changes in vascular network architecture. For LS174T tumors, non-labelled vessels (post-Oxi4503) were generally located in the centre of tumors, whereas SW1222 displayed a more distributed and localised pattern of perfusion loss (**Figure 4b**). Blood volume post-Oxi4503 significantly decreased, relative to baseline, in both tumour types (p<0.01, **Figure 4c**). In control tumours, no significant decrease was found in the degree of vessel labelling between baseline and 90 minutes post-saline (p>0.05).

### REANIMATE simulations of response to Oxi4503

REANIMATE simulations of steady-state fluid dynamics, based on vascular networks pre- and post-Oxi4503, revealed a spatially heterogeneous response, with both increases and decreases in perfusion and IFP observed within the same tumors, representing a redistribution of flow in response to localised vascular occlusion. These trends were replicated in *in vivo* data, which showed a significant decrease in median tumor perfusion of 9.8% (from 0.61 to 0.55 mL min^-1^ 100g^-1^, p<0.05) in LS174T tumors (**Figure 5**), but which was accompanied by a significant increase in the 90^th^ percentile perfusion value (from 2.48 to 2.64 mL min^-1^ 100g^-1^, p<0.01). Our simulations also predicted a decrease in interstitial fluid pressure (IFP) of 4.5 mm Hg, but was also accompanied by an increase of 3.6 mm Hg in the 90^th^ percentile. These results demonstrate a complex redistribution of flow caused by the vascular disrupting agent at this early time point. In SW1222 tumors, *in vivo* measurements of perfusion and IFP did not significantly change. Moreover, the fragmented nature of dual-labelled fluorescence images meant that post-Oxi4503 steady-state simulations could not be performed.

**Figure 5.**
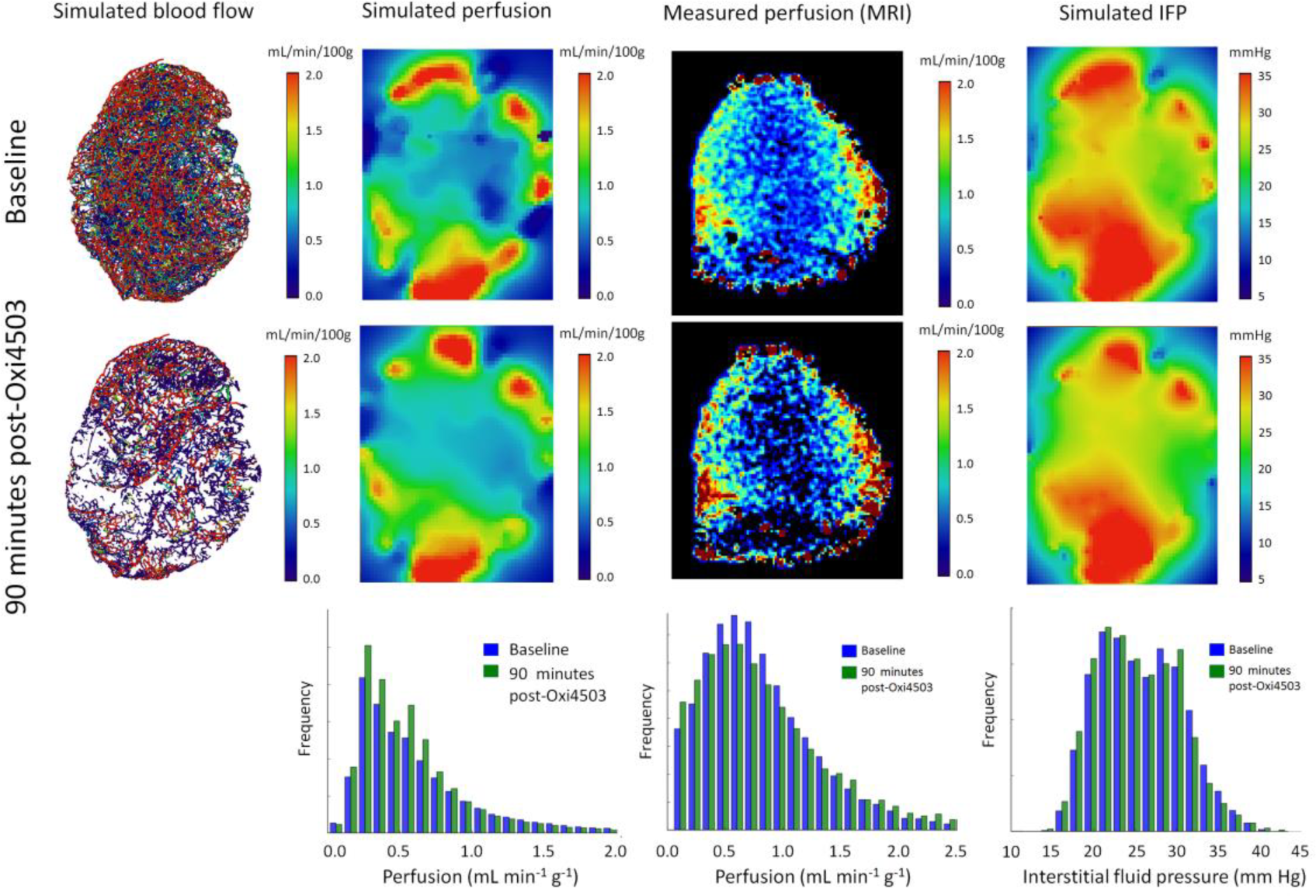
Results of REANIMATE simulations of blood flow, perfusion and interstitial fluid pressure (IFP), in an LS174T tumor, at baseline (top row) and 90 minutes post-Oxi4503 treatment (middle row). Images showing the change in perfusion *in vivo*, measured with arterial spin labelling MRI, are also shown, alongside histograms of each parameter (bottom row). Small changes in each parameter were observed, which were heterogeneously distributed throughout the tumor, both in simulations and *in vivo*.

We simulated the uptake of Oxi4503 as per Gd-DTPA simulations, but over a longer duration (90 minutes, with a temporal resolution of 10 seconds). Oxi4503 has a molar mass of 332.35 g mol^-1^ (approximately one third that of Gd-DTPA), so, using the Stoke-Einstein relation, *D* was set at 7.37×10^-5^ mm^2^ s^-1^. Systemic pharmacokinetics for Oxi4503 were taken from the literature, ^32^ and expressed as an exponential decay function (Equ. 15). Both intravascular and interstitial drug concentrations were recorded (see **Figure 6a**, **Supplemental Movie 4**), and the intravascular concentration used to measure the total blood vessel exposure to Oxi4503 (defined as the total amount of drug multiplied by the time exposed to it, per unit surface area of the vessel) (**Figure 6b**). As with Gd-DTPA experiments, Oxi4503 uptake was spatially heterogeneous.

**Figure 6.**
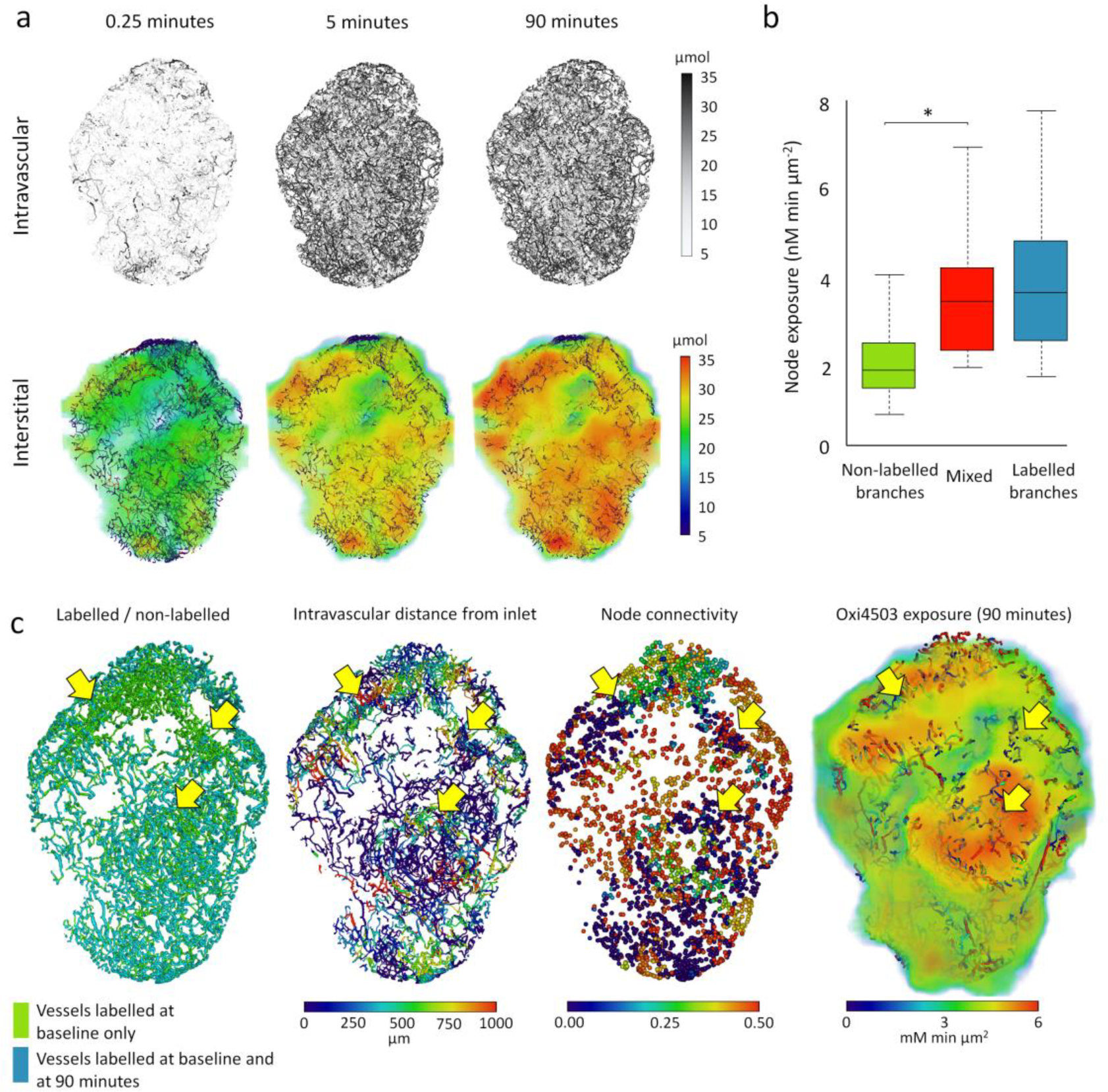
REANIMATE simulation predictions of Oxi4503 delivery and treatment response. (a) Maps of whole-tumor intravascular and interstitial (tissue) delivery of Oxi4503 from baseline to 90 minutes post-dosing. (b) Box plot of simulated Oxi4503 exposure in branch points connecting single-labelled only, dual-labelled only or a mixture of single- and dual-labelled vessels at 90 minutes post-Oxi4503 delivery. A significantly lower exposure to Oxi4503 was found in in single-labelled vessels. (c) A 1 mm slice through an LS174T vessel network, showing the location of dual- and single-labelled vessel segments, their intravascular distance from an inlet node, their connectivity score, and simulated Oxi4503 exposure (intravascular and interstitial). Arrows show the location of single-label clusters, which are associated with a larger intravascular distance from an inlet, lower node connectivity and mixed (both high and intermediate) Oxi4503 exposure.

These results show that, alongside realistic predictions from steady-state fluid dynamics simulations and Gd-DTPA uptake, REANIMATE can also provide realistic predictions of the response to the vascular targeting agent Oxi4503.

### Treatment-focussed hypothesis generation and testing with REANIMATE

We lastly used REANIMATE to test two hypotheses: 1) that vessels that receive the greatest Oxi4503 exposure are more likely to become non-perfused; 2) that network geometry differences between tumor types could influence their response to VDA therapy.

To test the first hypothesis, we compared vessels from our dual-labelled datasets that had lost perfusion post-Oxi4503 (i.e. were labelled with just one fluorphore), with their simulated exposure to Oxi4503, as predicted by REANIMATE. The result of this analysis was the opposite to that expected: that non-perfused vessels exhibited a signficantly lower simulated exposure than vessels that remained perfused (2.0 compared with 3.8 mM min m^-2^ (p<0.001)) (**Figure 6c**). This contradicts our first hypothesis, that perfusion loss would be associated with greater Oxi4503 dose, so was rejected. This lead us to next evaluate our second hypothesis, and investigate differences in the vascular architecture of the two tumour types. In particular, we evaluated the functional connectivity of the two tumor types.

Functional (or logical) connectivity and redundancy measures describe the connectedness of individual vessel networks, following pathways of decreasing fluid pressure. Specifically, redundancy was measured by *N*, the mean number of viable alternative pathways for each node if the shortest path (based on flow velocity) were occluded, and *r,* the average additional distance that would be travelled. Connectivity was defined as the sum of the number of nodes upstream and the number of nodes downstream of a given node, divided by the total number of nodes in the network.

This analysis showed us that SW1222 tumors have significantly greater vascular connectivity than LS174T tumors (*C* = 0.15 ± 0.06 and 0.06 ± 0.05, respectively) (p<0.01) (**Figure 2e**). They also display greater redundancy, with *N* = 1.9 ± 0.9 and 1.5 ± 0.7 and *r* = 1.02 ± 0.02 and 1.04 +/- 0.05, respectively (**Figure 2e**). We also found that nodes in LS174T vascular networks that connected both perfused and non-perfused vessels had signficantly greater predicted Oxi4503 exposure than branch points connecting non-perfused vessels (**Figure 6d**) (p<0.01).

Referring back to the definitions of connectivity and redundancy provided above, these results suggest that, in LS174T tumors, vessels that become non-perfused due to targeting by a high concentration of Oxi4503 can cause large vascular territories downstream to become nonperfused, due to their lack of connectivity. In SW1222 tumours, loss of perfusion can be compensated for by rerouting flow through alternative routes, thanks to their high redundancy. This explaination, which requires further evaluaiton, would explain the different pattern of response observed in SW1222 and LS174T tumors, with LS174T tumors showing large regions of perfusion loss in dual-fluorophore data (particularly in their core), whilst SW1222 show a more distributed pattern with flow loss in individual vessels.

## Discussion

We have presented REANIMATE, a novel, large-scale, three-dimensional imaging, modelling and analysis framework. REANIMATE uses advances in optical imaging of cleared tissue ^16,33,34^ to produce realistic substrates for computational modelling, and is guided by *in vivo* imaging data. We applied REANIMATE to imaging data from two murine models of colorectal cancer to simulate: 1) steady-state fluid dynamics (blood flow, intravascular and interstitial fluid pressure); 2) uptake of the MRI contrast agent Gd-DTPA; and 3) uptake and response to vascular-targeting treatment (Oxi4503). Our results demonstrate the feasibility and accuracy of this novel, whole-tissue approach to numerical modelling, which allows computational (*in silico*) experiments to be performed on real-world tumours.

Computational modelling of cancer has a relatively long history,^35-38^ and has allowed us to develop our understanding of, for example, tumour angiogenesis and microenvironmental structure in response to therapy, such as tumour blood flow,^39^ oxygen transport ^40^ and VEGF distribution.^41^ REANIMATE builds on and extends this work by including simulation substrates from complete, real-world tumors, in three spatial dimensions, which are guided by and compared against *in vivo* measurements. In this initial demonstration, we focussed on the vasculature of colorectal xenograft models, which we imaged by labelling with fluorescent lectin. This allowed blood flow to be explicitly simulated in realistic networks, providing new insights into drug delivery and blood flow heterogeneity. We treated the interstitium as a continuum, but future analyses could include additional structural elements such as cell membranes and nuclei, by multifluorescence labelling. Indeed, there is significant potential for extending and enhancing REANIMATE in other pathologies, to allow more in-depth computational experiments to be performed.

When we compared our REANIMATE predictions with *in vivo* imaging data, we found a good correspondence with the magnitude and spatial heterogeneity of *in vivo* measurements, both in steady-state and time-dependent simulation parameters. Perfusion was predicted to be signficantly greater in SW1222 tumors than in LS174T tumors, which reflects the results of *in vivo* arterial spin labelling measurements, and was highly heterogenous, with flow concentrated at the periphery of both tumor types. Conversely, Gd-DTPA uptake was more heterogeneous in LS174T than in SW1222 tumors, both in *in vivo* MRI measurements and simulations (with minimal parameter optimisation).

We focussed here on drug delivery, but REANIMATE could easily be modified to include response to therapy, for example by reducing interstitial cell density to reflect response to a cytotoxic agent. Tumors are normally spatially heterogeneous, and we have shown here that adding realistic, whole-tumour microstructure allows accurate predictions for drug delivery and tumor pathophysiology to be made. We showed that structural connectivity and redundancy in colorectal tumor xenograft model vascular networks can introduce different responses to the vascular-targeting agent Oxi4503. SW1222 tumors, with their greater connectivity and redundancy, are more able to resist loss of flow in individual vessels, by rerouting flow via local pathways, whereas there is much greater potential for LS174T tumors to lose perfusion in large downstream subnetworks.

Importantly, these results demonstrate a mechanism through which tumors can exhibit a form of physical resistance to drug therapies, which manifests via complex interactions across large regions within a tumor, or across whole organs. We have previously observed a similar heterogeneous response when assessing the response of colorectal metastases in the liver, in which ^9^ the magnitude of the response decreased with increasing distance of individual tumors from major blood vessels. This is again indicative of treatment resistance via physical mechanisms, such as vascular geometry and poor blood flow.

The detailed insights generated in this study could not have been made with conventional two-dimensional analysis of histological sections, or *in vivo* experiments that lack the spatial resolution and functional information to access this information, and demonstrates a key strength of the REANIMATE approach. We anticipate that REANIMATE will enable us to study and understand complex interactions between biological phenomena, allowing new insights into key challenges in cancer research. This could include a better understanding of limitations in tumor drug delivery (and how this can be mediated) ^36,42^ and the development of resistance to therapy via physical (rather than biochemical) mechanisms.^43^

## Materials and Methods

### Tumour xenograft models

All experiments were performed in accordance with the UK Home Office Animals Scientific Procedures Act 1986 and UK National Cancer Research Institute (NCRI) guidelines ^44^. 8-10 week old, female, immune-compromised nu/nu nude mice (background CD1) were used throughout this study (Charles River Laboratories). Human colorectal adenocarcinoma cell lines (SW1222 and LS147T) were cultured in complete media (Minimum Essential Medium Eagle with L-Glutamine (EMEM) (Lonza, Belgium) + 10% fetal bovine serum (Invitrogen, UK)) in a ratio 1:20 (v/v) and incubated at 37 °C and 5% CO_2_. To prepare for injection, cells were washed with DPBS and detached with trypsin-EDTA (7-8 min, 37 °C, 5% CO_2_). A 100 μl bolus of 5x10^6^ cells was injected subcutaneously into the right flank above the hind leg. Tumour growth was measured daily with callipers, for between 10 to 14 days.

### Fluorescent labelling of tumour vasculature and perfusion fixation

Lectin (griffonia simplicifolia) bound to either Alexa-647 (Thermo Fisher Scientific, L32451) or Alexa-568 (Thermo Fisher Scientific, L32458) was injected intravenously (i.v.) and allowed to circulate for 5 minutes, prior to perfuse fixation, to allow sufficient binding to the vascular endothelium.^3^

To prevent blood clot formation within the vasculature, mice were individually heparinized (Wockhardt) by intraperitoneal (i.p.) injection (0.2 ml, with 1000 IU ml^-1^). Mice were terminally anaesthetized by i.p. injection of 100 mg kg^-1^ sodium pentobarbital (Animalcare, Pentoject) diluted in 0.1 ml phosphate buffered saline (PBS). Once anaesthesia was confirmed, surgical procedures for intracardial perfusion were performed for systemic clearance of blood. PBS (30 ml, maintained at 37 °C) was administered with a perfusion pump (Watson Marlow, 5058) at a flow rate of 3 ml/min to mimic normal blood flow. After the complete drainage of blood, 40 ml of 4% paraformaldehyde (PFA, VWR chemicals) was administered. Harvested tumours were stored for 12 hours in 4% PFA (10 ml total volume, at 4 °C).

### Treatment with Oxi4503

Following 10 to 14 days of growth, mice were randomly assigned to treatment (Oxi4503, n=6) and control (saline) groups, with n=3 SW122 and n=3 LS174T in each. Treated groups were injected i.v. with 100 μg lectin-AlexFluor 647 diluted in sterile saline at neutral pH (100 μl) containing 1 mM CaCl_2_, followed by administration of OXi4503 (40 mg kg^-1^, 4 mg ml^-1^). Control mice were injected with 100 μl saline. After 2 hours, all mice were injected i.v. with 100 μg lectin-AlexFluor 568 diluted in sterile saline at neutral pH (100 μl) containing 1 mM CaCl_2_. 5 minutes after injection mice were culled and underwent perfuse-fixation, as described above.

### Optical clearing and imaging

Following perfuse-fixation, tumours were rinsed three times in PBS, for 10 minutes each, prior to clearing, to remove residual formaldehyde and avoid over-fixation.^45^ After PBS rinsing, harvested tumours were optically cleared with BABB (1:2 benzyl alcohol: benzyl benzoate). Our BABB clearing preparation consisted of dehydration in methanol for 48 hours followed by emersion in BABB for 48 hours.^16^

Fluorescently-labelled tumour vasculature in cleared tissue was visualized with optical projection tomography (OPT, Bioptonics, MRC Technologies, Edinburgh). Autofluorescence from tumour tissue was used to form an image of tumour morphology (excitation range 425/40 nm, emission range LP475 nm). Lectin-AlexaFluor 647 was imaged using a filter set with excitation range 620/60 nm, and emission 700/75 nm. For vessels labelled with lectin-AlexaFluor 568, a filter set with excitation 560/40 nm and emission LP610 nm was used. Measurements were performed with an exposure time of 1600-2000 ms for lectin-AlexaFluor 647 and of 270-600 ms for lectin-AlexaFluor 568, which was varied according to sample size. The rotation step was 0.45 degrees. The final resolution ranged from 4.3 μm to 8 μm, depending on the sample size.^46^

OPT data were reconstructed with Nrecon (Bruker, Ettlingen, Germany). Misalignment compensation was used to correct misalignment during projection image acquisition, in order to reduce tails, doubling or blurring in the reconstructed image. Depth of correction for ring artifact reduction was 4 and defect pixel masking was 50% for all scans.

### Image processing and vessel segmentation

Reconstructed OPT data were used to generate whole-tumor blood vessel networks. First, a three-dimensional Gaussian filter with a width of 50 pixels (corresponding to a physical size of 300 μm, greater than the largest vessel diameter) was applied. The filtered data were subtracted from the original data to remove background variations in autofluorescence. A three-dimensional Frangi filter was then applied (Matlab, MathWorks, Natick, MA) to enhance vessel-like structures.^17^ The response to the filter was thresholded to segment blood vessels from background. Skeletonisation of these thresholded data was performed in Amira (Thermo Fisher Scientific, Hillsboro, OR), which also converted the data into graph format (i.e. nodes and segments with associated radii). To ensure that vessel structures were accurately represented, 2D sections from the original image data were swept through reconstructed 3D networks (in Amira), with visual inspection used to for an accordance between vessel location and thickness, and the location of fluorescence signal (see **Supplemental Figure 2**).

### Mathematical model of steady-state tissue fluid dynamics

Blood flow through the segmented vascular network was modelled by Poiseuille’s law, using empirically-derived laws for blood viscosity (assuming constant network haematocrit) and following the established approach developed in ^47,48^. This model assumes conservation of flux at vessel junctions to define a linear system to solve for the pressures at nodal points in the network (from which vessel fluxes are calculated using Poiseuille). Boundary conditions on terminal nodes in the network were estimated using the optimisation method of Fry et al.,^49^ which matches the network solution to target mean shear stress and pressure values.

The approach of Fry et al.^49^ requires a proportion of boundary conditions to be applied to a microvascular network. However, neither flow or pressure measurements were obtained *in vivo* for our tumour networks. As such, a stochastic procedure was employed to induce a arbitrary pressure drop (55 to 15 mmHg for both LS147T and SW1222 simulations) across a network.^50^ In addition, consistent with previous studies,^51,52^ 33% of internal nodes were assigned zero flow. All remaining boundary nodes were unknowns. This procedure ensured physiologically realistic tissue perfusion when compared to that gathered *in vivo* using ASL MRI.

The network flow solution was coupled to an interstitial fluid transport model, adapting the approach taken in Secomb et al. ^53^ to model oxygen delivery to tissue. The interstitium is modelled as a porous medium using Darcy’s law,

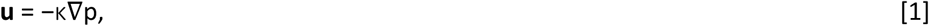

with ∇ · **u** = 0 and subject to *p* → *p_I_* as ***x*** → *∞*. Here, ***u*** is the volume-averaged interstitial blood velocity (IFV), *p* is the interstitial fluid pressure (IFP), *p_I_* is the target IFP, and κ is the conductance of the interstitial tissue. Starling’s law is used to describe fluid transport across the endothelium, from the vessels into the interstitium:

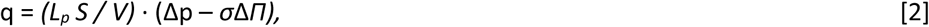

where, q is the fluid flux across the endothelium, *L_p_* is the hydraulic conductance of the vessel wall, *S* is the surface area of the vasculature, *V is* the tissue, *σ* is the oncotic reflection coefficient and, Δp and Δ*Π* fluid and oncotic pressure gradients between the vasculature and tissue.

To solve the model computationally, we discretised the tumor vasculature into a series of *N* sources of strength *q_sj_* so that the conservation of mass equation is modified to

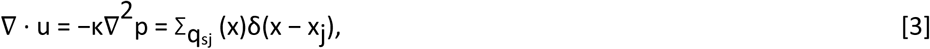

where ***x_j_*** and *q_sj_* are the spatial coordinates and (unknown) strength at ***x**_j_* of source *j*, respectively, and *δ(**x-x**_j_)* is the three-dimensional delta function. An axisymmetric Greens solution, *G(r)* where *r = |**x** – **x**_j_|*, was sought for equation 4 subject to the boundary condition that *p* → *p_I_* as ***x*** → *∞*.

Distributing the delta function *j* uniformly over a sphere of finite radius *r_0j_* (set to the radius of blood vessel *j*), the solution to equation 3 may be approximated by

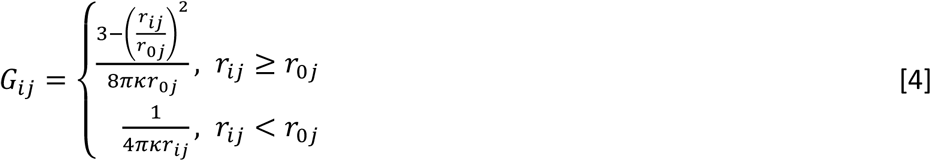

where *r_ij_ =|x_i_ – x_j_|* is the distance between sources *i,*j ϵ *N.* The corresponding interstitial fluid pressure (IFP) at source *i* may be approximated by

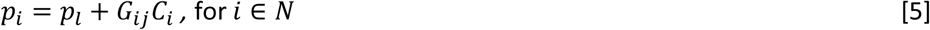

Assuming fluid flux across the vessel wall is continuous yields

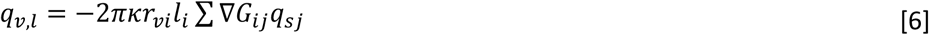

Assuming fluid pressure is continuous at the interface between the interstitial domain and the vasculature, Starling’s law can be written in the form

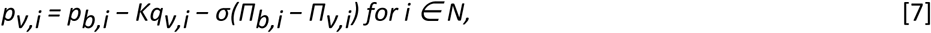

where *pv,i* and *Πv,i* are the blood and osmotic plasma pressure at the vessel wall, *pb,i* and *Πb,i* is the intravascular blood in the absence of diffusive interstitial fluid transfer (calculated using the Poiseuille flow model) and osmotic fluid pressure, *qv,i* is the rate of fluid flow per unit volume from blood vessel *i* to the interstitium. The intravascular resistance to fluid transport in blood vessel *i*, is defined by *K = V / L_p_ S.*

Combining equations 5, 6 and 7 can used to form a dense linear system, which can be solved for IFP. Full details on the mathematical model can be found in ^50^ with the corresponding parameter values used here shown in **Table 2**.

**Table 2.** Summary statistics for REANIMATE steady-state flow simulations in SW1222 and LS174T human colorectal carcinoma xenografts. Data are mean ± standard deviation, and P is the p-value associated with Wilcoxon rank sum tests to compare parameter means.

| Parameter (unit) | LS174T | SW1222 | P |
| --- | --- | --- | --- |
| Perfusion (mL min<sup>-1</sup> 100g<sup>-1</sup>) | 56 ± 7 | 73 ± 3 | <0.001 |
| Blood pressure (mm Hg) | 34.54 ± 2.07 | 34.83 ± 2.01 | >0.05 |
| Blood flow (nl min<sup>-1</sup>) | 21.9 ± 1.1 | 102.27 ± 11.3 | <0.001 |
| Blood velocity (mm s<sup>-1</sup>) | 0.83 ± 0.03 | 0.45 ± 0.04 | <0.001 |
| Shear stress dyn cm<sup>2</sup>) | 12.83 ± 0.43 | 6.19 ± 0.34 | <0.001 |
| Interstitial fluid pressure (mm Hg) | 33 ± 18 | 36 ± 12 | >0.05 |

### Mathematical model of time-dependent vascular and interstitial transport

A ‘propagating front’ (PF) algorithm was developed to describe the transport of solute (e.g. a drug) through the tumor vessel network and interstitium. This model considers the timescale for delivery of a drug, on which the flow problem is assumed to be steady (the timescales for drug transport by advection and diffusion are much faster than those for vascular adaption, which would contribute to a non-steady flow solution). A vascular input function was first defined, which describes the time-dependent delivery of the drug concentration into the network, and which then propagates throughout the network according to the network topology and flow solution. The influence of each vessel network inlet was modelled independently and each solution linearly superimposed, allowing the algorithm to be parallelised.

Each node was assigned a set of values, *J* describing the ratio of the flow in each vessel segment connected to the node (*F*), to the total inflow into the node (*F_in_*). Flow values were taken from the steady-state model defined above. *Q* values were propagated through the network, following pathways with decreasing vascular pressure. Using velocities from the steady-state solution, a set of delays, *d*, were also assigned to each node. Vessel segments attached to each node were catergorised as outflows (negative pressure gradient) or inflows (positive pressure gradient). Time-dependent drug concentration in the *k*^th^ outflowing vessel segment (*C*_k_(t)) was modelled as

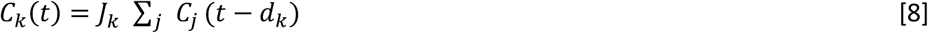

where *C_j_*(*t*) is the concentration in each inflowing vessel segment. Within vessel segments, vascular permeability to drug transport was modelled according to

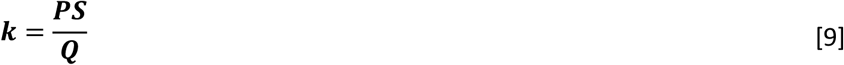

where *S* is the internal surface area of the vessel, *P* is the vessel wall permeability and *Q* is the blood flow (Poiseuille). *P* was initially fixed at 10^-4^ m s^-1^.^19,20^ Interstitial delivery was cast in a forward finite difference framework, in which vessels were considered as one-dimensional emitters. Points were gridded on concentric cylinders, regularly spaced around the vessel segment (with spacing ranging from 10 to 100 μm). Exchange of the drug across the vessel wall and diffusion through the interstitium were modelled as

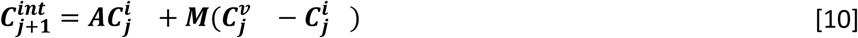

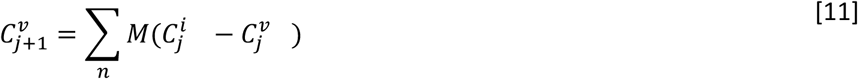

where 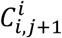 is the interstitial concentration at the *j^th^* time point and *C^v^_j_* is the vascular concentration. Interstitial velocity and pressure were not used in the time-dependent model, for simplicity, but could be incorporated in future studies, particularly for large molecules. *A* is a two-dimensional square matrix of dimension *n*, where *n* is the number of radial positions in the interstitial finite difference calculation, *h* is their radial separation and *k* is the spacing between time steps:

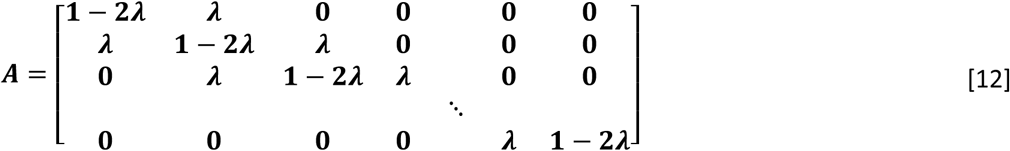

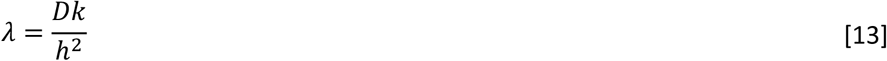

*D* is the diffusion coefficient of the agent under investigation. Following each finite difference step, interstitial diffusion solutions were regridded to a course 64×64×64 matrix (approximately 100 μm isotropic resolution) for storage. During regridding, absolute numbers of moles were converted to molar concentration.

### Measurement of vessel network functional connectivity and redundancy

The mean number of viable alternative pathways, *N*, for each node if the shortest path (based on transit time – i.e. incorporating flow velocity) was occluded was used to define the redundancy of tumor vessel networks, alongside *r,* the average additional distance that would be travelled.^54^ Connectivity was defined as the sum of the number of nodes upstream and the number of nodes downstream of a given node, divided by the total number of nodes in the network. All three measures reflect functional connectivity (i.e. following pathways with decreasing vascular pressure from steady-state fluid dynamics simulations), and were estimated from vessel networks using algorithms written in-house in Python 2.7.

### Simulation of Gd-DTPA delivery

The systemic pharmacokinetics for Gd-DTPA in mice, following an i.v. bolus injection, were modelled as a biexponential decay:

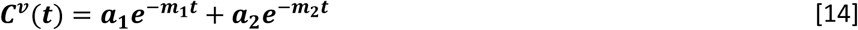

with *a*_1_ = 2.55 mM, *m*_1_ = 8×10^-2^ s^-1^, *a*_2_ = 1.2 mM and *m*_2_ = 1×10^-3^ s^-1^.^24^

### Simulation of Oxi4503 delivery

Oxi4503 systemic pharmacokinetics were modelled as a single exponential function, of the form

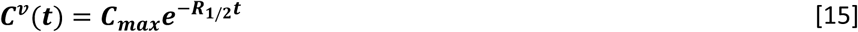

with *R*_1/2_ = 3.1×10^-5^ s^-1^ and *C*_max_ = 7.7 μM.^32^ This assumed a mouse mass of 25 g and injection dose of 40 mg kg^-1^.

### Dynamic contrast-enhanced MRI

Gadolinium-DTPA (Magnevist, Bayer, Leverkusen, Germany) was injected as a bolus into mouse tail veins, using a power injector (Harvard Instruments, Cambourne, UK). We injected 5 mL kg^-1^ over a period of 5 seconds, which was initiated at 90 seconds after the start of a dynamic, spoiled gradient-echo sequence (TE, 2.43 ms; TR, 15 ms; flip angle 20°; 5 slices; slice thickness 0.5 mm; matrix size, 128×128; FOV, 35×35 mm; temporal resolution 16 s; total duration 15 minutes). The change in signal intensity induced by contrast agent was calculated by subtracting the mean signal from the first 5 frames from the acquisition.

Signal intensity was converted to gadolinium concentration, via the change in longitudinal relaxation rate *R*_1_ and contrast agent relaxivity (*c*_1_ fixed at 2.9 mM^-1^ s^-1^):

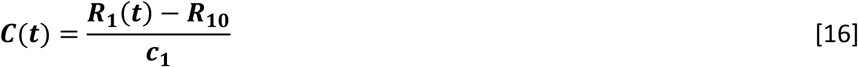

R1(*t*) was estimated from the theoretical change in spoiled gradient-echo signal magnitude.^55^ R10 was the mean, pre-enhancement *R*_1_, which was estimated from a Look-Locker multi-inversion time acqusition,^56^ acquired prior to the dyunamic sequence (TE, 1.18 ms; inversion time spacing, 110 ms; first inversion time, 2.3 ms; 50 inversion recovery readouts).

Contrast agent uptake data were fitted to a phenomenological model of the form

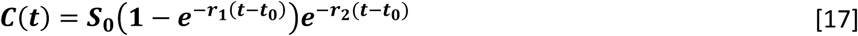

where *S*_0_, *r*_1_, *r*_2_ and *t*_0_ were fitted parameters. Fitting was performed in Python 2.7 (leastsq algorithm from the scipy package).

### Arterial spin labelling MRI

We acquired arterial spin labeling (ASL) data with a flow-sensitive alternating inversion recovery (FAIR) Look-Locker ASL sequence, with a single-slice spoiled gradient echo readout (echo time, 1.18 ms; inversion time spacing, 110 ms; first inversion time, 2.3 ms; 50 inversion recovery readouts; 4 averages).^9,56^ Regional perfusion maps were calculated as described by Belle et al. (38), with an assumed blood-partition constant of 0.9.

### Statistics

Differences between groups were tested for significance with the non-parametric Wilcoxon rank sum test (Python 2.7, scikit package). P < 0.05 was considered significant. All summary data are presented as mean ± SD.

## Acknowledgements

We acknowledge the support received for the Kings College London & UCL CR-UK and EPSRC Comprehensive Cancer Imaging Centre, in association with the MRC and Department of Health (England), (C1519/A10331), Wellcome Trust (WT100247MA), Rosetrees Trust / Stoneygate Trust (M135-F1 and M601). We thank OXiGENE for supplying Oxi4503.

